# Genome Sequence of Segmented Filamentous Bacteria Present in the Human Intestine

**DOI:** 10.1101/813196

**Authors:** Hans Jonsson, Luisa W. Hugerth, John Sundh, Anders F. Andersson

**Affiliations:** Department of Molecular Sciences, Swedish University of Agricultural Sciences, 75007 Uppsala, Sweden; Center for Translational Microbiome Research, Department of Molecular, Tumour and Cell Biology, Karolinska Institutet, 17177 Stockholm, Sweden; Stockholm University, Department of Biochemistry and Biophysics, National Bioinformatics Infrastructure Sweden, Science for Life Laboratory, 17121 Stockholm, Sweden; Science for Life Laboratory, Department of Gene Technology, KTH Royal Institute of Technology, 17121 Stockholm, Sweden

**Keywords:** Segmented filamentous bacteria, SFB, intestinal microbiota, metagenome, metagenome-assembled genome, human intestine, host-bacterial interaction, immune modulation

## Abstract

Segmented filamentous bacteria (SFB) colonize the small intestine of a variety of animals in a host-specific manner. SFB are physically attached to the host’s intestinal epithelium and affect several functions related to the immune system, among them IgA production and T-cell maturation. Until now, no human-specific SFB genome had been described. Here, we report the metagenomic reconstruction of an SFB genome from a human ileostomy sample. Phylogenomic analysis clusters the genome with the SFB genomes from mouse, rat and turkey, but the genome is genetically distinct, displaying 65-71% average amino acid identity to the other genomes, and is tentatively unique for the human small intestine. By screening human faecal metagenomic datasets, we identified individuals carrying sequences identical to the new SFB-genome. We thus conclude that a unique SFB variant exists in humans and we foresee a renewed interest in the elucidation of SFB functionality in this environment.

## Introduction

The interdependency of the intestinal microbiota and its host manifests itself in various ways, of which some on the host side are highly spectacular. Thus, effects spanning from improving nutrient uptake (Krajmalnik-Brown et al., 2012) or metabolizing drugs (Zimmermann et al., 2019) to influencing risk of cancer (Feng et al., 2015) and altering cognitive function (Martin et al., 2018) have been reported to be dependent on microbiota composition and functionality. Although this area of research has attracted great attention during the last decades, the codes for communication between microbes and man have only begun to be deciphered.

Investigations of intestinal host-microbe interactions have been revolutionized by the development and application of powerful DNA sequencing and bioinformatics tools. Nevertheless, although huge amount of data is gained in this way, the translation of these data into a meaningful context lingers. Another, somewhat interlinked approach, has been to search for tentative “key players” which share an evolutionary history with the host and maintain important host functions through specific mechanisms. The identification of such key players is not straightforward and although some exquisite examples exist (Atarashi et al., 2013, 2015; Tan et al., 2016; Tanoue et al., 2019), only a limited number of commensal bacteria has so far been identified as having defined effects on their host. Segmented filamentous bacteria, SFB, represents one of these key players and holds a so far unique capacity to elicit full maturation of the mouse gut immune barrier. The work with SFB during the last decades beautifully describes the cross-fertilization between different areas of research, particularly microbiology and immunology (Cerf-Bensussan, 2019).

SFB were discovered already in the mid-1960s in laboratory animals (Hampton and Rosario, 1965; Savage, 1969) where they could be identified by microscopy due to their filamentous growth and the unique attachment of the filament to the intestinal wall. Several intriguing features connected to the lifestyle of these organisms later opened for a deeper interest in their possible role as important symbionts (Davis and Savage, 1974). Thus, they colonized primarily in the terminal part of the small intestine where many immune cells are located; they appeared at greater number around weaning which is an important period for maturation of the immune system, and, not least, they exhibited an intimate contact with the host through a specific anchoring to the intestinal cell wall. Together, these observations led to speculations and later also the first reports that SFB affected immune functions of the host (Klaasen et al., 1993; Umesaki et al., 1995). After these early observations, SFB have been subject to a large number of studies (reviewed in (Ericsson et al., 2014; Schnupf et al., 2017)) which has firmly established their role as immunomodulatory bacteria. Thus, they are attributed with a long row of effects, including stimulation of chemokine and antimicrobial components production, induction of gut lymphoid tissue and a strong increase in faecal IgA (Umesaki et al., 1999). However, their potent triggering of T helper 17 (T_H_17) cell differentiation is perhaps their most eye-catching attribute (Ivanov et al., 2009) in terms of immunomodulation. Interestingly, very recent experiments applying immunodeficient mice, demonstrated the ability of SFB to also confer protection against rotavirus infection independent of immune cells (Shi et al., 2019).

Although not yet cultivable, SFB mono-colonized laboratory animals have offered a route to isolation and characterization of this group of organisms. Complete genomes are available from SFB isolated from mice (Bolotin et al., 2014; Kuwahara et al., 2011; Prakash et al., 2011; Sczesnak et al., 2011) and rats (Prakash et al., 2011) and an unpublished draft genome sequence from turkey is publicly available at NCBI (GenBank accession number GCA_001655775.1). Genomic analysis has revealed that SFB are gram-positive spore-forming bacteria with a distinct phylogenetic position within the Clostridales. They have small genomes of around 1.5 Mb which is reflected by a limited biosynthetic capability, rendering them a functional position between free-living bacteria and obligate intracellular symbionts (Bolotin et al., 2014; Sczesnak et al., 2011).

A hallmark of SFB biology seems to be host specificity, as supported both by experimental and genetic data. Thus, colonization experiments have shown that cross-colonization with mouse SFB in rats or rat SFB in mice is not possible (Tannock et al., 1984). This argues for different species or lineages in SFB adapted to different hosts. After the discovery of SFB in rodents and with the accumulating evidence for their ability to affect crucial steps in immune development, it was natural to search for them also in humans. The first study indicating their presence in humans visualized a tentative SFB organism adherent to ileal biopsied tissue by light microscopy (Klaasen et al., 1993). More recently, 16S rRNA gene sequences of SFB were reported in human samples using SFB-specific PCR primers; Yin et al. (Yin et al., 2013) found SFB sequences in 55 faecal samples while one of us (Jonsson) detected an SFB sequence in an ileostomy sample (Jonsson, 2013). While the faecal SFB sequences were phylogenetically interleaved with SFB sequences from mice from the same study, the ileostomy sequence was distinct from SFB sequences from other animals. Except for the 16S sequences from these studies, no human SFB DNA sequences have been published. Moreover, human gut shotgun metagenomic sample sets have been scanned by attempting to map reads to the SFB genomes of mice and rats, but without success (Sczesnak et al., 2011). Thus, up until now, no genomic data have been presented for human-derived SFB, and it is still an open question whether a human-adapted variant of the organism actually exists.

We now report the draft genome sequence of a tentatively human-adapted representative of the SFB group. With metagenomic approaches, we have reconstructed the SFB genome from the same ileostomy sample that earlier produced the unique 16S rRNA gene sequence. Phylogenetic analysis clusters this genome to the SFB genomes described earlier, yet clearly defines it as unique. In addition, we could show the presence of sequences derived from the new genome in unrelated individuals through screening of published metagenome data. Our data strengthen the likelihood that the paradigm with host-specific colonization is valid also for SFB-human symbiosis. Considering the possibility of analogous immune-modulatory activities of SFB in humans and rodents, this finding could be of paramount importance.

## Results and Discussion

### Genome reconstruction

To verify the presence of an SFB 16S rRNA gene sequence in the human ileostomy sample where it was earlier detected with SFB-specific primers, we subjected the same sample to amplicon sequencing using broad-taxonomic range PCR primers. This confirmed the existence of an SFB sequence: after sequence noise removal, a single amplified sequence variant (ASV) was classified as *Candidatus* Arthromitus and this was identical over its full length to the previously published 16S sequence from the same sample. The relative abundance of this ASV was however low, as it represented 0.16 - 0.37% of the microbial community’s ASV sequences, depending on the DNA extraction method used.

In order to assemble the genome of the candidate SFB organism, we conducted deep shotgun metagenomic sequencing using Illumina NovaSeq, which generated a total of 953,167,834 read-pairs for four different DNA preparations from the same sample. The 317,687 contigs of the resulting assembly were binned into genomes using information on sequence composition and coverage. To improve the binning procedure, the coverage of the contigs was estimated not only using the four different DNA libraries from the sample that were prepared using three different DNA extraction methods, but also using publicly available human gut metagenomes. We tried two different binning software, CONCOCT and MetaBAT2, and applied two different contig length cutoffs for each binning software. The two binners generated approximately the same number of bins with comparable quality estimates (Figure S1), but only MetaBAT2 generated a bin at each length cutoff that was classified as SFB (genus *Savagella* according to the Genome Taxonomy Database (GTDB)). These two bins differed by a few contigs, and we used a conservative approach of defining the SFB metagenome-assembled genome (MAG) as all contigs shared by both bins (127 contigs, 1,221,164 bp), as well as those uniquely found in one but taxonomically classified as SFB (*Candidatus* Arthromitus according to NCBI; 25 contigs, 89,165 bp). As is often the case for MAGs, a contig encoding a 16S rRNA gene was missing. rRNA gene prediction however identified a 4.2 kb contig encoding a 16S gene with a region identical to our SFB amplicon sequence, and this contig could be linked to contigs of the MAG using read-pair information. The contig (k141_89555) encodes a full-length 16S gene as well as a 23S gene. The 16S gene is 96% similar across its full length to those encoded in the SFB mouse and rat genomes. Notebly, it has mismatches to commonly used primers for PCR identification of SFB (Figure S2). Adding this contig resulted in a 1,314,549 bp (153 contig) MAG, that we denote SFB-human-IMAG (IMAG; ileostomy metagenome-assembled genome). SFB-human-IMAG has a single-copy gene-estimated completeness and contamination of 85.6% and 0%, respectively (Table S1).

### Phylogenomic analysis

The reconstructed genome was subjected to phylogenomic analysis using a set of universally conserved protein sequences. This verified the placement of SFB-human-IMAG among the SFB (with 100% support). Intriguingly, the human-assembled SFB genome was most closely related to SFB isolated from turkey (GCA_001655775.1) and the two formed a sister clade to the SFB genomes from mouse and rat (Figure 1). This pattern was supported by average amino acid identity (AAI) analysis, with SFB-human-IMAG displaying 71% AAI to SFB-turkey, while displaying 65% AAI to the SFB from rodents (Table 1). It was however not supported by a phylogenetic tree based solely on the full-length 16S genes of the genomes (Figure S3). The conflicting phylogenies between the SFB and their hosts could indicate that the SFB have switched hosts during the course of evolution. It could also reflect that the human and turkey SFB belong to a different lineage than the mouse and rat SFB, and that the two lineages diverged before mammals diverged from birds. The two SFB lineages may exist in all hosts, or one could have gone extinct in some of the hosts.

**Table 1.**
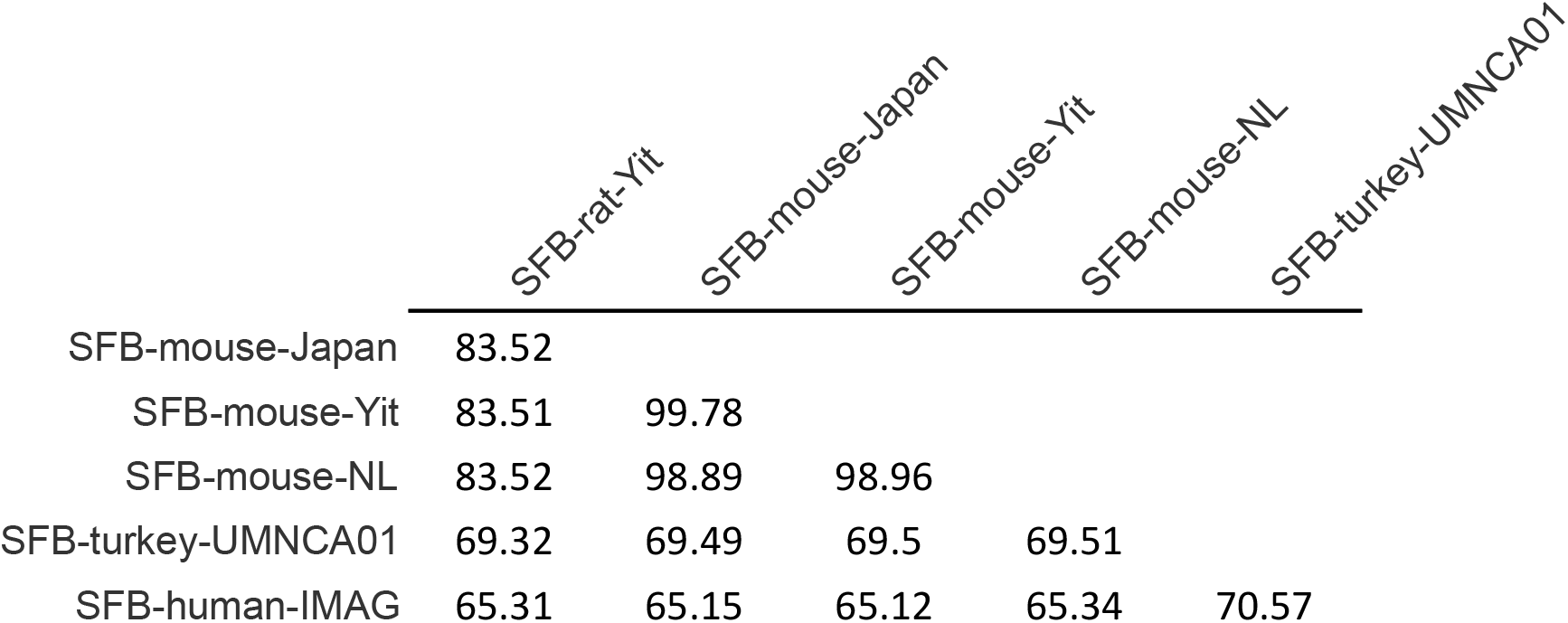
Average amino acid identity between sequenced SFB genomes.

|  | SFB-rat-Yit | SFB-mouse-Japan | SFB-mouse-Yit | SFB-mouse-NL | SFB-turkey-UMNCA01 |
| --- | --- | --- | --- | --- | --- |
| SFB-mouse-Japan | 83.52 |  |  |  |  |
| SFB-mouse-Yit | 83.51 | 99.78 |  |  |  |
| SFB-mouse-NL | 83.52 | 98.89 | 98.96 |  |  |
| SFB-turkey-UMNCA01 | 69.32 | 69.49 | 69.5 | 69.51 |  |
| SFB-human-IMAG | 65.31 | 65.15 | 65.12 | 65.34 | 70.57 |

**Figure 1.**
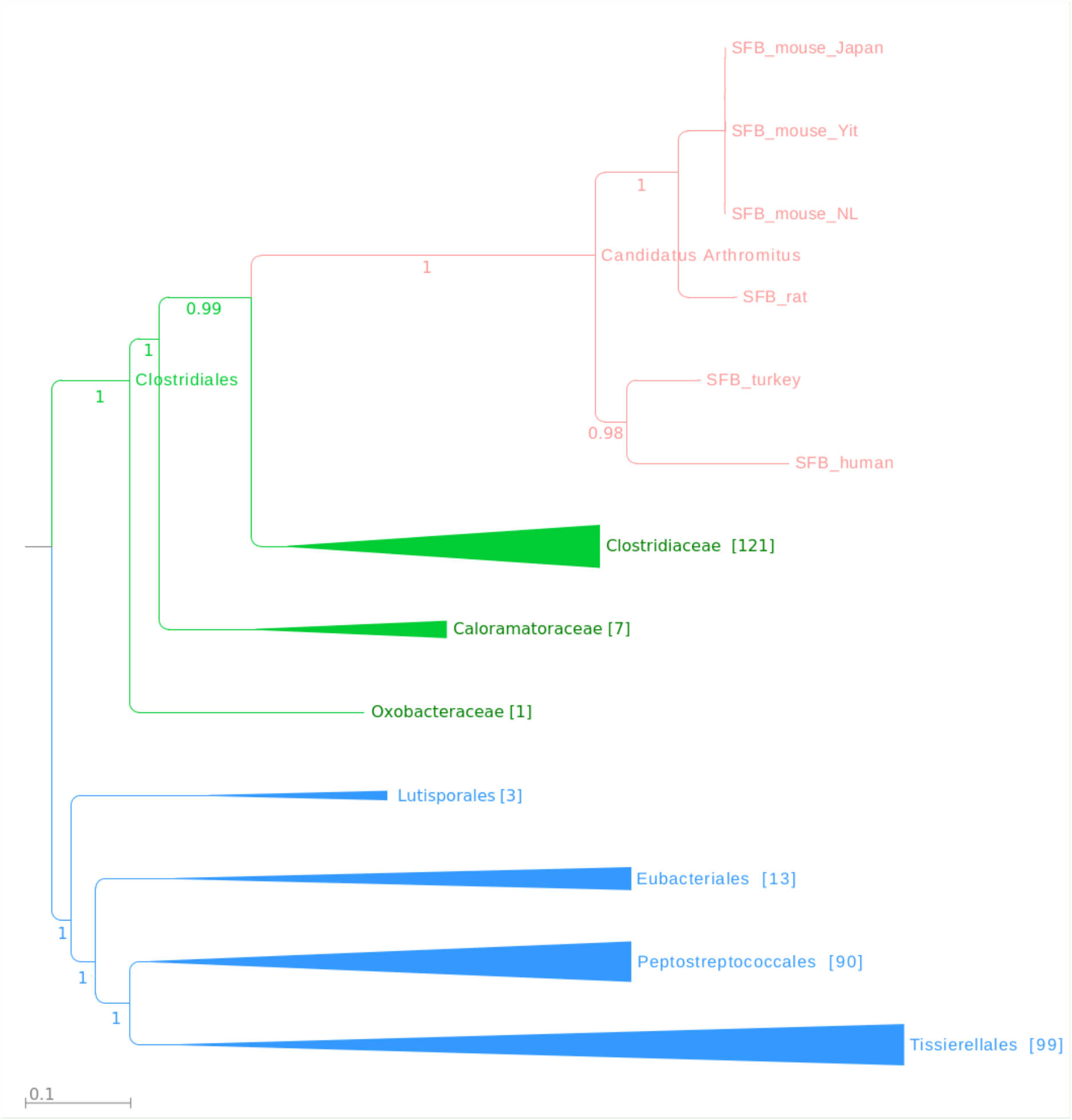
Phylogenomic tree of genome-sequenced SFB and related Clostridia. Internal branches are marked with support values (range 0 - 1). The genus *Candidatus* Arthromitus is highlighted in red; other families of order Clostridiales are depicted in green. Orders that form a monophyletic sister group to Clostridiales are shown in blue.

### Gene content and physiology

SFB-human-IMAG encodes 1276 proteins of which 1117, 1018 and 798 could be assigned to eggNOG/COG, PFAM and KO families, respectively. The percentage of genes assigned to different eggNOG functional categories did not differ markedly between SFB-human-IMAG and the other SFB genomes (Figure S4), indicating that the organisms have similar functional capabilities. Nevertheless, 29 of the COGs found in SFB-human-IMAG were missing in all the rodent SFB (Figure 2; Table S2); twelve of which were also found in SFB-turkey. Several of the 29 COGs, like DNA methylase, transfer proteins TraG and TraE and SNF2 family helicases, appear to be encoded on prophage sequences or other mobile elements. Conversely, SFB-human-IMAG misses 114 COGs that were found in all the other SFB, but most of these are likely missing due to that the genome is incomplete.

**Figure 2.**
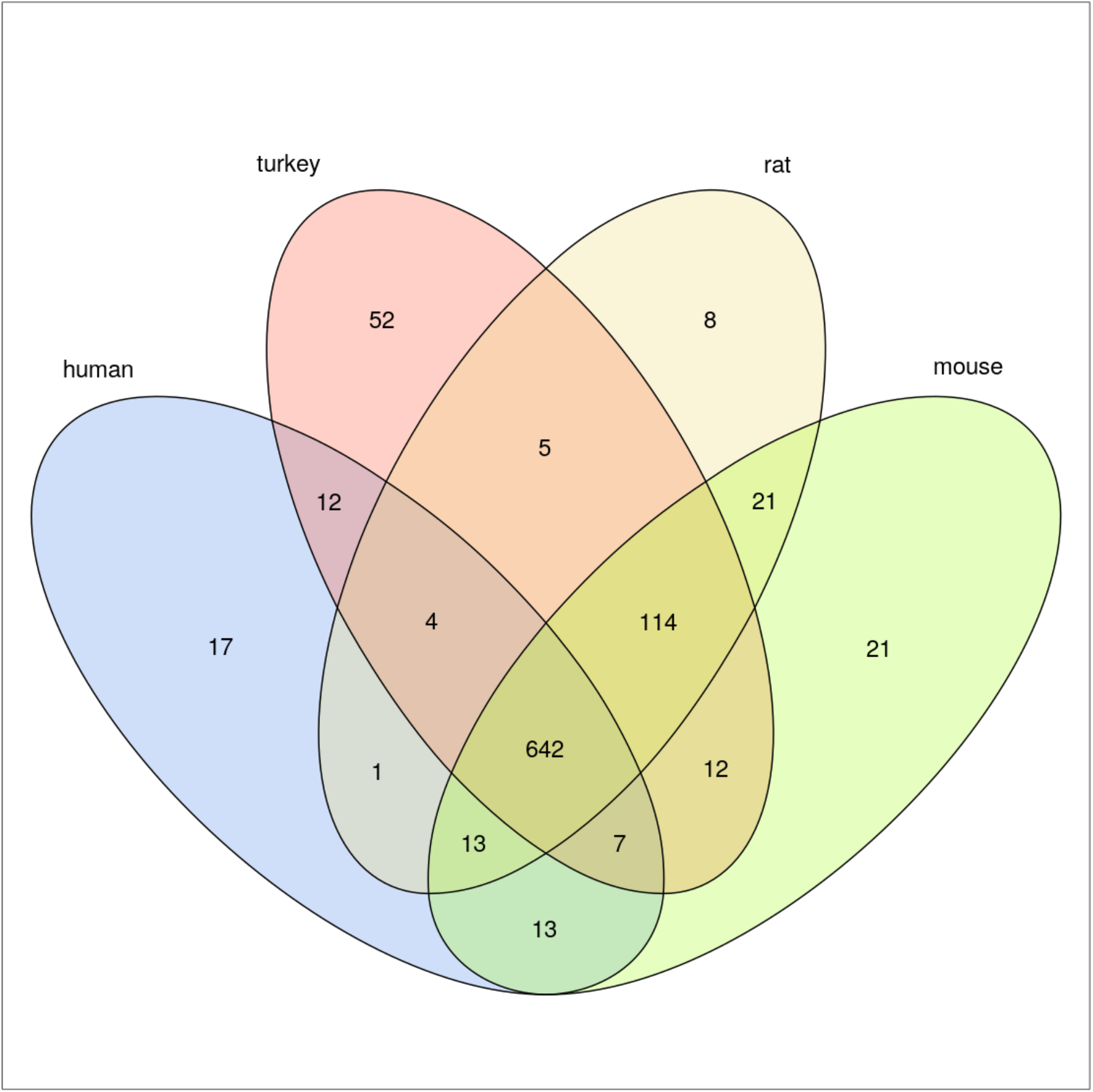
Venn diagram depicting the overlap in COG content for the SFB lineages from different hosts. The mouse lineage represents the union of SFB-mouse-Yit, SFB-mouse-Japan and SFB-mouse-NL, since only 24 COGS are not shared amongst all of these.

SFB has previously been described as having a fermentative metabolism. We have identified all but a few of the enzymes for glucose utilization also in the draft genome of SFB-human-IMAG. No enzymes involved in the tricarboxylic acid cycle were identified, and, accordingly, there are no proteins that can be assumed to take part in an electron transport chain, confirming a fermentative lifestyle. As previously described SFB, SFB-human-IMAG appears to have a restricted capability to synthesize amino acids, vitamins/cofactors and nucleotides. One interesting observation, however, is that SFB-human-IMAG contains six genes for biotin synthesis (BIOA, B, D, F, W and X), suggesting it has the capability to synthesize biotin. The corresponding genes show sequence homology to genes from SFB-turkey but most of them are lacking in the SFB from rodents. This could reflect differences in the physiology or diet of the hosts, or differences in the microbial community in which rodent and human SFB exist, since some gut microbes can synthesise this cofactor while others are auxotrophs. A limited biosynthesis machinery inevitably results in a dependency on the host environment for nutrient supply. Nine glycoside hydrolases (GH) representing six different GH families (Table S3), one tentatively extracellular N-acetylglucosaminidase, and several cell surface-bound and extracellular proteases were identified in the draft genome. Together, these enzymes are likely used for harvesting components from the intestinal milieu.

The SFB-human-IMAG genome also encodes a large number of transport functions. This is in agreement with a restricted metabolic capability and similar to other SFB. Since the genome is not complete, some transport functions are likely missing due to incomplete genome assembly. A notable exception is the lack of the ABC transporter for phosphonate, where the specific genes are missing in the middle of an SFB-human-IMAG contig that otherwise displays conserved synteny with SFB-mouse-Japan. However, since phosphorous is indispensable, bacteria have evolved several systems for acquisition of this macronutrient, and SFB, including SFB-human-IMAG, carry genes for a phosphate specific transport system (sfb.merged_01113 - sfb.merged_01115).

When comparing with the annotation of the complete genome of SFB-mouse-Japan, we conclude that SFB-human-IMAG is likely to carry a complete set of genes for sporulation and germination. Likewise, a complete set of genes for flagellar motility and chemotaxis are present, and it is thus reasonable to assume that the bacterium has the ability for motility and chemotaxis.

### Host-microbe interactions

The intestinal microbiota influences the host locally or systemically through a number of mechanisms. These could be of an indirect character such as the production of vitamins or other metabolites that interact with host functions. The intestinal microbiota as a whole is responsible for such effects, and thus, specific interactions and individual microbial contributions may be concealed. More direct effects could be mediated by bacteria that physically interact with host structures. While the bulk of our knowledge about such interactions comes from studies of various pathogens, the increasing knowledge of SFB biology has the potential to change this. These bacteria interact with their host in a way not seen with other commensal/symbiotic bacteria and a number of proteins and functions of SFB have been proposed as instrumental in host association and immune modulatory effects. Among these are immunogenic flagellins, tentative fibronectin binding proteins which could effectuate binding to host cell matrix, phospholipase C and ADP-ribosyltransferase, both influencing actin polymerization which is a characteristic at sites of SFB attachment, and others (Kuwahara et al., 2011; Pamp et al., 2012; Prakash et al., 2011; Sczesnak et al., 2011). Due to the prevailing limitations in culture and genetic manipulation, however, very limited experimental data pinpointing the role of individual SFB components in host interactions are available. One example though, is the ability of SFB flagellins to interact *in vitro* with TLR5 receptors and to activate the NF-kB signalling pathway and elicit the innate immune response(Kuwahara et al., 2011). We identified three flagellins in the genome of SFB-human-IMAG and at least two of these contain a conserved motif for TLR5 recognition and activation (Song et al., 2017)(Figure S5).

SFB interacts physically with intestinal enterocytes through polar attachment of the SFB-filament and triggers an invagination of the enterocyte without breaching the cell membrane. This intimate contact suggests a strong potential for interaction with the host, and indeed, data were recently published that show how SFB in mice can transfer cell wall proteins into the enterocyte (Ladinsky et al., 2019). This protein (p3340) was earlier shown to be a major target in the antigen-specific CD4 T_H_17 cell response induced by SFB(Yang et al., 2014). The corresponding protein is also encoded in the SFB-human-IMAG genome (sfb.merged_00774). It is interesting to note that while the N-terminal (signal sequence) and the C-terminal parts of these proteins display high amino acid identity, the main part shows only a low degree of identity (Figure S6). In the work by Yang et al. (Yang et al., 2014) two peptides from p3340 were reported to strongly stimulate T_H_17 cells. These peptides are conserved only to a limited degree in SFB-human-IMAG, leaving open the possibility that the variability in sequence reflects host adaptation and thus the evolvement of human-specific T_H_17 triggering epitopes.

The components of SFB responsible for attachment to the enterocytes have not been identified. Secreted and cell surface located bacterial proteins generally play major roles in signal transduction, ion transport and host cell adhesion. While enzymes and transporters often contains signature motifs, many proteins involved in adhesion are undefined as to their functional sites, and therefore depicted as hypothetical. A number of secreted and cell surface proteins were predicted in our genome based on N-terminal signal peptides and 60 of these are hypothetical proteins. The size of the hypothetical and tentatively extracellular proteins in SFB-human-IMAG ranges from 57 to 2040 amino acids, and the identity to homologous proteins from SFB from other animal hosts are in the range of 34-72%, with a mean of 52% identity. This is substantially lower than the overall identity of the SFB-Human-IMAG proteome with other SFB (Table 1), and thus indicates more rapid evolution in proteins communicating with the exterior environment. It is plausible that some of these proteins play a role in attachment and host communication and thereby mediate the host-specificity that is a characteristic of SFB.

SFB do not harbor a gene for sortase, the enzyme that normally anchors many cell surface proteins in gram-positive bacteria, and a corresponding mechanism has not been described in the SFB-group. It is likely though that an alternative route for anchoring of cell surface proteins exists in SFB. Interestingly, a conserved amino acid motif located C-terminally was earlier identified in a number of putative cell surface proteins in SFB(Pamp et al., 2012). We have localized this motif in a number of predicted extracellular proteins, including the T_H_17 stimulating protein p3340 from mouse SFB and the orthologous protein 00774 from SFB-Human-IMAG mentioned above. Supporting evidence for anchoring comes from the study of Ladinsky et al (Ladinsky et al., 2019), where immuno EM shows the location of p3340 to the SFB cell wall. We therefore postulate that SFB-human-IMAG has at least 14 cell surface proteins that may be anchored to the bacterial surface through the involvement of this aa-motif. Furthermore, twelve proteins encoded by the SFB-Human-IMAG genome are predicted to be anchored via a lipoprotein motif, and six proteins could possibly be anchored via an N-terminal transmembrane helix (TMHMM 2). This leaves a substantial number of predicted extracellular proteins seemingly anchorless. While some of these likely are true secretory proteins, it is notable that a number of them have a very high isoelectric point, giving them a basic charge which in turn could allow them to re-associate with the bacterial surface (Turner et. al 1997).

### Presence in other metagenomes

To verify that SFB-human-IMAG resides in the human intestine, we searched for it among published metagenomes from the human gut. The genome was first BLAST-searched against an integrated catalogue of reference genes in the human gut microbiome (IGC 9.9) (Chen et al., 2019), which consists of 9.9 million genes assembled from 1,267 human faecal samples. Only fourteen of the IGC genes gave matches to the genome when requiring ≥95% identity and ≥70% of the IGC gene’s bases aligned. However, this gene catalogue is mainly derived from samples from adults, while SFB in most animals peak in young individuals during weaning (Jiang et al., 2001). Therefore, we instead scanned a large recent metagenomic study consisting of a time-series of faecal samples from children 0 - 3 years born in Russia, Estonia and Finland (Vatanen et al., 2016). The reads from the metagenome samples were first mapped against SFB-human-IMAG using standard settings. This rendered substantial mapping for many samples. However, manual inspection of the alignments revealed that the mapped reads were typically only partially aligned, and to regions displaying unusually high sequence conservation, such as structural RNA genes. Redoing the mapping with stringent settings (see Methods) and only counting reads mapped to protein-coding genes (CDS) gave substantially reduced mapping; however, 61 out of the 153 contigs were mapped by at least one read pair, and 7 out of the 817 samples had at least one read-pair mapping. Two of these samples, one Estonian infant at day 390 (SRS1719092) and one Finnish infant at day 320 (SRS1719390), had particularly many reads mapping and mapped with 1-3 read pairs each to 24 and 44 contigs, respectively. Although the SFB-mapping reads only corresponded to three and eleven out of a million mapped reads, respectively, in these samples, the reads appeared to be randomly distributed over the genome, indicating that the genome is present in these samples, rather than that some genome regions are wrongly binned or horizontally transferred. In comparison, zero reads from the 817 samples mapped to any of the rodent SFB genomes using the same settings, and the draft SFB genome from turkey was mapped only at two CDS, both corresponding to genes being 100% identical at the nucleotide level to genes of several Firmicutes genomes. Since the infant metagenomes analysed were derived from another lab, it can be excluded that the mappings to SFB-human-IMAG are due to contamination of DNA from our Ileostomy sample or sequencing library. We also checked for the presence of SFB in the metagenomes from intestinal luminal fluids from three Chinese children that had earlier been screened positive for SFB with PCR (Chen et al., 2018). With the exception for reads mapping to one of the above SFB-turkey CDS, no mapping to any of the SFB genomes were obtained for these samples. In summary, our analyses show that SFB-human-IMAG is present in human infant faecal material, although in very low relative abundance.

## Conclusions

SFB holds a so far unique position in our collective knowledge on how individual components in the intestinal microbiota can affect host functions. The intimate interaction with the intestinal cells represents a remarkable evolutionary mechanism and recent data has shown that this is indeed a route for SFB-host interaction. Although SFB has been described from many host species, conclusive data regarding a human-specific SFB has been lacking. The data presented in this study strongly suggests that such a lineage actually exists. The assembled genome clusters with the previously described SFB genomes while being clearly distinct from these. The insight that SFB could be a natural component of the human microbiota calls for deepened attempts to elucidate their impact on human physiology in general and immune development in particular.

## Methods

### Sample collection and storage

Samples were initially collected and processed as described by (Lundin et al., 2004). Briefly, 10 adult subjects previously proctocolectomised for ulcerative colitis volunteered to participate in the experiment (two female subjects, eight males, age range 24–65 y, BMI 20.7-35.6 kg/m2). The subjects were living a normal life based on physical examination and blood tests before the experiment. The study was approved by the Ethical Committee of the Umeå University Hospital. Ileostomy bags were immediately frozen on dry ice and stored at −30°C. Ileostomy effluents from each 24h period were freeze-dried to constant weight, mixed, homogenized and stored at −70°C until analysis. One of the subjects was earlier (Jonsson 2013) identified as positive for the presence of an SFB-related 16S rRNA sequence on the basis of PCR analysis and sequencing. Sample from this individual was used in this work.

### DNA extraction

The DNA used for the 16S amplicon sequencing was extracted using QIAamp DNA Stool Mini Kit (QIAGEN, Venlo, Netherlands) with an added bead beating treatment as the first step. Bead beating was performed with 0.1 mm zirconium/silica beads (Biospec Products, Bartlesville, OK, USA), 2 x 45 s with setting 5 using the MP FastPrep□24 (MP Biomedicals, Irvine, CA, USA). Of the five samples used for amplicon sequencing, the first two were extracted from the original material while the latter three corresponded to three size fractions, selected by gravity precipitation. Since the SFB content of these size fractions was not significantly larger than for the full sample, all later DNA extractions were performed on unfractionated material. For the shotgun sequencing, two replicates were extracted with QIAamp DNA Stool Mini Kit and two with QIAamp Fast DNA Stool Mini Kit, and one additional replicate with QIAamp DNA Microbiome Kit, according to instructions from the manufacturer (QIAGEN, Venlo, Netherlands).

### 16S rRNA gene amplification and sequencing

DNA extracts were amplified using universal 16S primers 341f and 805r (Herlemann et al., 2011) enhanced with Illumina adapters as described by (Hugerth et al., 2014) (341f: 5′-ACACTCTTTCCCTACACGACGCTCTTCCGATCT-N5-CCTACGGGNGGCWGCAG-3′; 805r: 5’-AGACGTGTGCTCTTCCGATCTGGACTACHVGGGTWTCTAAT-3’, where N5 represents 5 random bases used to improve sequencing quality) using 25 µL of Kapa Hifi mastermix (Kapa Biosystems, Woburn, MA, USA), 2.5 µl of each primer (10 µM), 2.5 µl of template DNA (1 ng/µl), and 17.5 µl of water. These mixtures were submitted to thermocycling in a Mastercycler Pro S (Eppendorf, Hamburg, Germany) under the following conditions: 95°C for 5 min, 98°C for 1 min, 20 cycles of 98°C for 20 s, 51°C for 20 s, and 72°C for 12 s, followed by a final elongation step of 72°C for 1 min. The products of these reactions were cleaned as described by Lundin et al (Lundin et al., 2010), concentrating the product to 23 µL. These were then barcoded in a PCR reaction containing 25 µL Kapa Mastermix polymerase and 1 µL of each barcoding primer (5′-AATGATACGGCGACCACCGAGATCTACAC-X8-ACACTCTTTCCCTACACGACG-3 and 5′-CAAGCAGAAGACGGCATACGAGAT-X8-GTGACTGGAGTTCAGACGTGTGCTCTTCCGATCT-3′, where X8 is a barcoding sequence) and with the following cycling conditions: 95°C for 5 min, 98°C for 1 min, 10 cycles of 98°C for 10 s, 62°C for 30 s, and 72°C for 15 s, followed by a final elongation step of 1 min. The products were cleaned again as described by Lundin *et al.* and concentrated to 15 µL. DNA concentration was measured with Qubit dsDNA HS (Thermo Fisher Scientific, Waltham, MA, USA) and the length and purity of the amplified product was verified with BioAnalyzer 2100 DNA1000 (Agilent Technologies, Santa Clara, CA, USA).

The products were sequenced on Illumina MiSeq with 2×300 bp together with amplicon samples from a different project. Cutadapt v.1.18 (Martin, 2012) was used to remove primer sequences, 3’-bases with a Phred score <15, and sequences not containing the expected primers. The resulting sequences were submitted to Unoise3 (Edgar, 2016). Taxonomic annotation was performed with SINA based on SILVA 132 (Pruesse et al., 2012).

### Metagenomic library preparation and sequencing

Libraries were prepared with the ThruPLEX DNA-seq kit (Rubicon genomics, Ann Arbor, MI, USA), aiming at an average fragment length of 350 bp. Sequencing was performed in a NovaSeq 6000 in S1 mode, yielding 358-410 million reads/sample.

### Preprocessing of shotgun reads

For the ileostomy samples, adapters were trimmed from the sequences using cutadapt (Martin, 2012) (v. 1.18) with default settings using the adapter sequences AGATCGGAAGAGCACACGTCTGAACTCCAGTCAC (ADAPTER_FWD) and AGATCGGAAGAGCGTCGTGTAGGGAAAGAGTGTAGATCTCGGTGGTCGCCGTATCATT (ADAPTER_REV). Removal of *phiX* sequences was performed by aligning reads against the *phiX* genome (GCF_000819615.1) using bowtie2 (Langmead and Salzberg, 2012) (v. 2.3.4.3) with parameters ‘--very-sensitive’ and only keeping pairs that did not align concordantly. Duplicates were removed using fastuniq (Xu et al., 2012) (v. 1.1) with default settings. This was followed by a second cutadapt trimming step using parameters ‘-e 0.3 --minimum-length 31’. Reads were then classified taxonomically using kraken2 (Wood et al., 2019) (v. 2.0.7_beta). Reads classified as human were removed prior to assembly.

Three external datasets of human gut samples were used for binning and for checking the presence of the obtained SFB MAG: 21 samples from BioProject PRJNA288044 (unpublished), 785 samples from BioProject PRJNA290380 (Vatanen et al., 2018), and 11 samples from BioProject PRJNA299342 (Chen et al., 2018). The 21 PRJNA288044 samples and the 11 PRJNA299342 samples were preprocessed by adapter and quality trimming using Trimmomatic (Bolger et al., 2014) (v. 0.38) with parameters ‘PE 2:30:15 LEADING:3 TRAILING:3 SLIDINGWINDOW:4:15 MINLEN:31’ followed by removal of phiX sequences as above. The 785 PRJNA290380 samples were preprocessed in the same manner but with the NexteraPE adapters and with duplicate removal following the phiX filtering step.

### Assembly and binning

Preprocessed ileostomy shotgun reads were assembled using megahit (Li et al., 2015) (v. 1.1.3) with settings ‘--min-contig-len 300 --prune-level 3 --k-list 21,29,39,59,79,99,119,141’. Preprocessed reads from all samples (ileostomy and external samples) were then aligned against the assembled contigs using bowtie2 (Langmead and Salzberg, 2012) (v. 2.3.4.3) with ‘--very-sensitive’ settings, followed by duplicate removal using MarkDuplicates (picard v. 2.18.21 (Wysoker et al., 2010)) with default settings. This output was used to calculate contig abundance profiles in all samples using the jgi_summarize_bam_contig_depths script from metabat2 (Kang et al., 2019) (v. 2.12.1). Binning of assembled contigs was then performed in two runs using metabat2 with parameters ‘--seed 123 -m <min_contig_length>’ where ‘min_contig_length’ was set to 1500 and 2500 for the two runs. For binning using CONCOCT (Alneberg et al., 2014) (v. 1.0.0), contig abundance profiles were computed using the concoct_coverage_table.py script followed by binning in two separate runs, both using default settings but with minimum contig length (‘-l’) set to 1000 and 2500, respectively. Bin quality was assessed using checkm (Parks et al., 2015) (v. 1.0.13) using lineage-specific marker genes. Ribosomal RNA genes were identified on assembled contigs using barrnap (https://github.com/tseemann/barrnap) (v. 0.9) with parameters ‘--reject 0.1’.

### Taxonomic annotation of contigs

Assembled contigs were classified taxonomically using package tango (https://github.com/johnne/tango, v. 0.5.6) and the UniRef100 protein database (release 2019_02). The package queried contigs in a blastx search using diamond (Buchfink et al., 2015) (v. 0.9.22) with parameters ‘--top 5 --evalue 0.001’. From the results, contigs were assigned a lowest common ancestor from hits with bitscores within 5% of the best hit. Assignments were first attempted at species level using only hits at ≥ 85% identity. If no hits were available at that cutoff, an attempt was made to assign taxonomy at the genus level using hits at ≥ 60% identity, followed by the phylum level at ≥45% identity. These rank-specific thresholds were chosen from (Luo et al., 2014).

### Functional annotation of genome

The SFB-human-IMAG bin as well as five sequenced genomes of SFB (RefSeq accessions GCF_000284435.1 GCF_000709435.1 GCF_000283555.1 GCF_001655775.1 GCF_000270205.1) were annotated using prokka (Seemann, 2014) (v. 1.13.3) with default settings. The prokka pipeline includes tRNA identification with aragorn (v. 1.2.38), prediction of ribosomal RNA with barrnap (https://github.com/tseemann/barrnap) (v. 0.9), gene calling with prodigal (Hyatt et al., 2010) (v. 2.6.3), homology searching with blastp (Camacho et al., 2009)(v. 2.7.1+) and HMM-profile searches with hmmer (Finn et al., 2011)(v.3.2.1). Protein sequences predicted with prokka were further annotated using eggnog-mapper (Huerta-Cepas et al., 2017) (v. 1.0.3) in ‘diamond’ run mode with the 4.5.1 version of the eggNOG database. Kegg orthologs, enzymes, pathways and modules were inferred from the eggnog-mapper output using the Kegg brite hierarchy information. Proteins were also annotated with PFAM protein families using pfam_scan.pl (v. 1.6) with default settings and v. 31 of the PFAM database. Carbohydrate-active enzyme Annotations were inferred using hmmscan against the dbCAN (http://bcb.unl.edu/dbCAN/) database (v. 6), followed by parsing of the output with the hmmscan-parser.sh script downloaded from the dbCAN server (http://bcb.unl.edu/dbCAN/download/hmmscan-parser.sh) and filtering using settings recommended for bacteria in the dbCAN readme (E-value < 1e-18 and coverage > 0.35).

### Phylogenetic and amino acid similarity analyses

The phylogeny of the SFB genomes was inferred using GTDB-TK (Parks et al., 2018) (v. 0.2.2) with GTDB release86, in both ‘classify_wf’ and ‘denovo_wf’ modes. The former placed the query genomes into an existing reference tree using pplacer (Matsen et al., 2010) while keeping the reference tree intact and was used to assign a GTDB taxonomy to the genomes. The latter instead created a new phylogenetic tree using both reference and query genomes and was used to investigate the phylogenetic relationship between the genomes. In the ‘denovo_wf’ method FastTree (Price et al., 2010) (v. 2.1.10) was used with the WAG protein model and Gamma20-based likelihoods (‘-wag -gamma’).

For the 16S phylogenetic analysis, one full-length 16S rRNA gene from each of the previously published complete SFB genomes, as well as from the genomes of five different species of Clostridium, were downloaded from the RDP (Cole et al., 2014). The positioning of the 16S rRNA gene in SFB-human-IMAG contig k141_89555 and in SFB-turkey contig GCF.001655775_NZ_LXFF01000001.1 was predicted with CheckM. The six 16S genes were aligned with Muscle (Edgar, 2004) and columns with gaps removed with DegePrime (Hugerth et al., 2014). A phylogenetic tree was constructed with FastTree using the GTR+CAT modell (results were nearly identical using the Jukes-Cantor + CAT model).

Average amino acid identity (AAI) between genome pairs were calculated using the online AAI calculator (http://enve-omics.ce.gatech.edu/aai/index), using default parameter settings.

### Prediction of extracellular proteins

SignalP-5.0 was used to identify signal peptides in the translated ORFs of the SFB-human-IMAG draft genome. The setting of organism group was gram-positive.

### Quantifying SFB in external metagenomes

Matching of the ORFs in IGC v9.9 (db.cngb.org/microbiome) against SFB-human-IMAG was performed with blastn v2.7.1+ (Camacho et al., 2009) requiring at least 80% identity over at least 70% of the query sequence. To assess the presence of SFB-human-IMAG and of SFB from mouse, rat and turkey in the faeces of young children, we used the recent work of Vatanen *et al (Vatanen et al., 2016)*, one of the datasets that we used for the binning. Mapping of the preprocessed reads against the SFB genomes was run in ‘strict’ mode, where only alignments without mismatches were reported (‘--score-min C,0,0’ in bowtie2). Counts of read-pairs mapping inside protein-coding regions (CDS) was obtained with featureCounts(Liao et al., 2014) (v. 1.6.4) with settings ‘-p -B -M’ to only count read-pairs with both ends mapped and allowing multimapping reads. The same procedure was used for mapping the shotgun reads from Chen *et al* (Chen et al., 2018).

### Data and Code Availability

The preprocessed amplicon and shotgun sequencing reads generated during this study, and the contig sequences of SFB-human-IMAG bin, are available at the European Nucleotide Archive (ENA) under the study accession number PRJEB34939. Data files for amplicon sequence variants, genome annotations, phylogenomic analysis, genome quality estimates and metagenome read mappings are available at XXXX.

## Acknowledgements

DNA sequencing was conducted at the Swedish National Genomics Infrastructure (NGI) and at Clinical Genomics at Science for Life Laboratory (SciLifeLab) in Stockholm. Computations were performed on resources provided by the Swedish National Infrastructure for Computing (SNIC) through the Uppsala Multidisciplinary Center for Advanced Computational Science (UPPMAX). We are grateful to Lars Engstrand, CTMR/KI, for his contribution to shogun sequencing discussions.

## References

Alneberg, J., Bjarnason, B.S., de Bruijn, I., Schirmer, M., Quick, J., Ijaz, U.Z., Lahti, L., Loman, N.J., Andersson, A.F., and Quince, C. (2014). Binning metagenomic contigs by coverage and composition. Nat. Methods 11, 1144–1146.

Atarashi, K., Tanoue, T., Oshima, K., Suda, W., Nagano, Y., Nishikawa, H., Fukuda, S., Saito, T., Narushima, S., Hase, K., et al. (2013). Treg induction by a rationally selected mixture of Clostridia strains from the human microbiota. Nature 500, 232–236.

Atarashi, K., Tanoue, T., Ando, M., Kamada, N., Nagano, Y., Narushima, S., Suda, W., Imaoka, A., Setoyama, H., Nagamori, T., et al. (2015). Th17 Cell Induction by Adhesion of Microbes to Intestinal Epithelial Cells. Cell 163, 367–380.

Bolger, A.M., Lohse, M., and Usadel, B. (2014). Trimmomatic: a flexible trimmer for Illumina sequence data. Bioinformatics 30, 2114–2120.

Bolotin, A., de Wouters, T., Schnupf, P., Bouchier, C., Loux, V., Rhimi, M., Jamet, A., Dervyn, R., Boudebbouze, S., Blottière, H.M., et al. (2014). Genome Sequence of “Candidatus Arthromitus” sp. Strain SFB-Mouse-NL, a Commensal Bacterium with a Key Role in Postnatal Maturation of Gut Immune Functions. Genome Announc. 2.

Buchfink, B., Xie, C., and Huson, D.H. (2015). Fast and sensitive protein alignment using DIAMOND. Nat. Methods 12, 59–60.

Camacho, C., Colouris, G., Avagyan, V., Ma, N., Papadopoulos, J., Bealer, K., and Madden, T.L. (2009). BLAST+: architecture and applications. BMC Bioinformatics 10, 421.

Cerf-Bensussan, N. (2019). Microbiology and immunology: An ideal partnership for a tango at the gut surface-A tribute to Philippe Sansonetti. Cell. Microbiol. e13097.

Chen, B., Chen, H., Shu, X., Yin, Y., Li, J., Qin, J., Chen, L., Peng, K., Xu, F., Gu, W., et al. (2018). Presence of Segmented Filamentous Bacteria in Human Children and Its Potential Role in the Modulation of Human Gut Immunity. Front. Microbiol. 9, 1403.

Chen, I.-M.A., Chu, K., Palaniappan, K., Pillay, M., Ratner, A., Huang, J., Huntemann, M., Varghese, N., White, J.R., Seshadri, R., et al. (2019). IMG/M v.5.0: an integrated data management and comparative analysis system for microbial genomes and microbiomes. Nucleic Acids Res. 47, D666–D677.

Cole, J.R., Wang, Q., Fish, J.A., Chai, B., McGarrell, D.M., Sun, Y., Brown, C.T., Porras-Alfaro, A., Kuske, C.R., and Tiedje, J.M. (2014). Ribosomal Database Project: data and tools for high throughput rRNA analysis. Nucleic Acids Res. 42, D633–D642.

Davis, C.P., and Savage, D.C. (1974). Habitat, succession, attachment, and morphology of segmented, filamentous microbes indigenous to the murine gastrointestinal tract. Infect. Immun. 10, 948–956.

Edgar, R.C. (2004). MUSCLE: multiple sequence alignment with high accuracy and high throughput. Nucleic Acids Res. 32, 1792–1797.

Edgar, R.C. (2016). UNOISE2: improved error-correction for Illumina 16S and ITS amplicon sequencing.

Ericsson, A.C., Hagan, C.E., Davis, D.J., and Franklin, C.L. (2014). Segmented filamentous bacteria: commensal microbes with potential effects on research. Comp. Med. 64, 90–98.

Feng, Q., Liang, S., Jia, H., Stadlmayr, A., Tang, L., Lan, Z., Zhang, D., Xia, H., Xu, X., Jie, Z., et al. (2015). Gut microbiome development along the colorectal adenoma-carcinoma sequence. Nat. Commun. 6, 6528.

Finn, R.D., Clements, J., and Eddy, S.R. (2011). HMMER web server: interactive sequence similarity searching. Nucleic Acids Res. doi: 10.1093/nar/gkr367.

Hampton, J.C., and Rosario, B. (1965). The attachment of microorganisms to epithelial cells in the distal ileum of the mouse. Lab. Invest. 14, 1464–1481.

Herlemann, D.P.R., Labrenz, M., Jurgens, K., Bertilsson, S., Waniek, J.J., and Andersson, A.F. (2011). Transitions in bacterial communities along the 2000 km salinity gradient of the Baltic Sea. ISME J. 5, 1571–1579.

Huerta-Cepas, J., Forslund, K., Coelho, L.P., Szklarczyk, D., Jensen, L.J., von Mering, C., and Bork, P. (2017). Fast Genome-Wide Functional Annotation through Orthology Assignment by eggNOG-Mapper. Mol. Biol. Evol. 34, 2115–2122.

Hugerth, L.W., Wefer, H.A., Lundin, S., Jakobsson, H.E., Lindberg, M., Rodin, S., Engstrand, L., and Andersson, A.F. (2014). DegePrime, a program for degenerate primer design for broad-taxonomic-range PCR in microbial ecology studies. Appl. Environ. Microbiol. 80, 5116–5123.

Hyatt, D., Chen, G.-L., Locascio, P.F., Land, M.L., Larimer, F.W., and Hauser, L.J. (2010). Prodigal: prokaryotic gene recognition and translation initiation site identification. BMC Bioinformatics 11, 119.

Ivanov, I.I., Atarashi, K., Manel, N., Brodie, E.L., Shima, T., Karaoz, U., Wei, D., Goldfarb, K.C., Santee, C.A., Lynch, S.V., et al. (2009). Induction of intestinal Th17 cells by segmented filamentous bacteria. Cell 139, 485–498.

Jiang, H.Q., Bos, N.A., and Cebra, J.J. (2001). Timing, localization, and persistence of colonization by segmented filamentous bacteria in the neonatal mouse gut depend on immune status of mothers and pups. Infect. Immun. 69, 3611–3617.

Jonsson, H. (2013). Segmented filamentous bacteria in human ileostomy samples after high-fiber intake. FEMS Microbiol. Lett. 342, 24–29.

Kang, D., Li, F., Kirton, E.S., Thomas, A., Egan, R.S., An, H., and Wang, Z. (2019). MetaBAT 2: an adaptive binning algorithm for robust and efficient genome reconstruction from metagenome assemblies (PeerJ Preprints).

Klaasen, H.L., Koopman, J.P., Van den Brink, M.E., Bakker, M.H., Poelma, F.G., and Beynen, A.C. (1993). Intestinal, segmented, filamentous bacteria in a wide range of vertebrate species. Lab. Anim. 27, 141–150.

Krajmalnik-Brown, R., Ilhan, Z.-E., Kang, D.-W., and DiBaise, J.K. (2012). Effects of gut microbes on nutrient absorption and energy regulation. Nutr. Clin. Pract. 27, 201–214.

Kuwahara, T., Ogura, Y., Oshima, K., Kurokawa, K., Ooka, T., Hirakawa, H., Itoh, T., Nakayama-Imaohji, H., Ichimura, M., Itoh, K., et al. (2011). The lifestyle of the segmented filamentous bacterium: a non-culturable gut-associated immunostimulating microbe inferred by whole-genome sequencing. DNA Res. 18, 291–303.

Ladinsky, M.S., Araujo, L.P., Zhang, X., Veltri, J., Galan-Diez, M., Soualhi, S., Lee, C., Irie, K., Pinker, E.Y., Narushima, S., et al. (2019). Endocytosis of commensal antigens by intestinal epithelial cells regulates mucosal T cell homeostasis. Science 363.

Langmead, B., and Salzberg, S.L. (2012). Fast gapped-read alignment with Bowtie 2. Nat. Methods 9, 357–359.

Li, D., Liu, C.-M., Luo, R., Sadakane, K., and Lam, T.-W. (2015). MEGAHIT: an ultra-fast single-node solution for large and complex metagenomics assembly via succinct de Bruijn graph. Bioinformatics 31, 1674–1676.

Liao, Y., Smyth, G.K., and Shi, W. (2014). featureCounts: an efficient general purpose program for assigning sequence reads to genomic features. Bioinformatics 30, 923–930.

Lundin, E.A., Zhang, J.X., Lairon, D., Tidehag, P., Aman, P., Adlercreutz, H., and Hallmans, G. (2004). Effects of meal frequency and high-fibre rye-bread diet on glucose and lipid metabolism and ileal excretion of energy and sterols in ileostomy subjects. Eur. J. Clin. Nutr. 58, 1410–1419.

Lundin, S., Stranneheim, H., Petterson, E., Klevebring, D., and Lundeberg, J. (2010). Increased Throughput by Parallelization of Library Preparation for Massive Sequencing. PLoS One 5, e10029.

Luo, C., Rodriguez-R, L.M., and Konstantinidis, K.T. (2014). MyTaxa: an advanced taxonomic classifier for genomic and metagenomic sequences. Nucleic Acids Res. 42, e73.

Martin, M. (2012). Cutadapt removes adapter sequences from high-throughput sequencing reads. Bioinformatics in Action 17, 10–12.

Martin, C.R., Osadchiy, V., Kalani, A., and Mayer, E.A. (2018). The Brain-Gut-Microbiome Axis. Cell Mol Gastroenterol Hepatol 6, 133–148.

Matsen, F.A., Kodner, R.B., and Armbrust, E.V. (2010). pplacer: linear time maximum-likelihood and Bayesian phylogenetic placement of sequences onto a fixed reference tree. BMC Bioinformatics 11, 538.

Pamp, S.J., Harrington, E.D., Quake, S.R., Relman, D.A., and Blainey, P.C. (2012). Single-cell sequencing provides clues about the host interactions of segmented filamentous bacteria (SFB). Genome Res. 22, 1107–1119.

Parks, D.H., Imelfort, M., Skennerton, C.T., Hugenholtz, P., and Tyson, G.W. (2015). CheckM: assessing the quality of microbial genomes recovered from isolates, single cells, and metagenomes. Genome Res. 25, 1043–1055.

Parks, D.H., Chuvochina, M., Waite, D.W., Rinke, C., Skarshewski, A., Chaumeil, P.-A., and Hugenholtz, P. (2018). A standardized bacterial taxonomy based on genome phylogeny substantially revises the tree of life. Nat. Biotechnol. 36, 996–1004.

Prakash, T., Oshima, K., Morita, H., Fukuda, S., Imaoka, A., Kumar, N., Sharma, V.K., Kim, S.-W., Takahashi, M., Saitou, N., et al. (2011). Complete genome sequences of rat and mouse segmented filamentous bacteria, a potent inducer of th17 cell differentiation. Cell Host Microbe 10, 273–284.

Price, M.N., Dehal, P.S., and Arkin, A.P. (2010). FastTree 2--approximately maximum-likelihood trees for large alignments. PLoS One 5, e9490.

Pruesse, E., Peplies, J., and Glöckner, F.O. (2012). SINA: accurate high-throughput multiple sequence alignment of ribosomal RNA genes. Bioinformatics 28, 1823–1829.

Savage, D.C. (1969). Localization of certain indigenous microorganisms on the ileal villi of rats. J. Bacteriol. 97, 1505–1506.

Schnupf, P., Gaboriau-Routhiau, V., Sansonetti, P.J., and Cerf-Bensussan, N. (2017). Segmented filamentous bacteria, Th17 inducers and helpers in a hostile world. Curr. Opin. Microbiol. 35, 100–109.

Sczesnak, A., Segata, N., Qin, X., Gevers, D., Petrosino, J.F., Huttenhower, C., Littman, D.R., and Ivanov, I.I. (2011). The genome of th17 cell-inducing segmented filamentous bacteria reveals extensive auxotrophy and adaptations to the intestinal environment. Cell Host Microbe 10, 260–272.

Seemann, T. (2014). Prokka: rapid prokaryotic genome annotation. Bioinformatics 30, 2068–2069.

Shi, Z., Zou, J., Zhang, Z., Zhao, X., Noriega, J., Zhang, B., Zhao, C., Ingle, H., Bittinger, K., Mattei, L.M., et al. (2019). Segmented Filamentous Bacteria Prevent and Cure Rotavirus Infection. Cell.

Song, W.S., Jeon, Y.J., Namgung, B., Hong, M., and Yoon, S.-I. (2017). A conserved TLR5 binding and activation hot spot on flagellin. Sci. Rep. 7, 40878.

Tan, T.G., Sefik, E., Geva-Zatorsky, N., Kua, L., Naskar, D., Teng, F., Pasman, L., Ortiz-Lopez, A., Jupp, R., Wu, H.-J.J., et al. (2016). Identifying species of symbiont bacteria from the human gut that, alone, can induce intestinal Th17 cells in mice. Proc. Natl. Acad. Sci. U. S. A. 113, E8141–E8150.

Tannock, G.W., Miller, J.R., and Savage, D.C. (1984). Host specificity of filamentous, segmented microorganisms adherent to the small bowel epithelium in mice and rats. Appl. Environ. Microbiol. 47, 441–442.

Tanoue, T., Morita, S., Plichta, D.R., Skelly, A.N., Suda, W., Sugiura, Y., Narushima, S., Vlamakis, H., Motoo, I., Sugita, K., et al. (2019). A defined commensal consortium elicits CD8 T cells and anti-cancer immunity. Nature 565, 600–605.

Umesaki, Y., Okada, Y., Matsumoto, S., Imaoka, A., and Setoyama, H. (1995). Segmented filamentous bacteria are indigenous intestinal bacteria that activate intraepithelial lymphocytes and induce MHC class II molecules and fucosyl asialo GM1 glycolipids on the small intestinal epithelial cells in the ex-germ-free mouse. Microbiol. Immunol. 39, 555–562.

Umesaki, Y., Setoyama, H., Matsumoto, S., Imaoka, A., and Itoh, K. (1999). Differential roles of segmented filamentous bacteria and clostridia in development of the intestinal immune system. Infect. Immun. 67, 3504–3511.

Vatanen, T., Kostic, A.D., d’Hennezel, E., Siljander, H., Franzosa, E.A., Yassour, M., Kolde, R., Vlamakis, H., Arthur, T.D., Hämäläinen, A.-M., et al. (2016). Variation in Microbiome LPS Immunogenicity Contributes to Autoimmunity in Humans. Cell 165, 842–853.

Vatanen, T., Franzosa, E.A., Schwager, R., Tripathi, S., Arthur, T.D., Vehik, K., Lernmark, Å., Hagopian, W.A., Rewers, M.J., She, J.-X., et al. (2018). The human gut microbiome in early-onset type 1 diabetes from the TEDDY study. Nature 562, 589–594.

Wood, D.E., Lu, J., and Langmead, B. (2019). Improved metagenomic analysis with Kraken 2.

Wysoker, A., Tibbetts, K., McCowan, M., Homer, N., and Fennell, T. (2010). Picard Tools.

Xu, H., Luo, X., Qian, J., Pang, X., Song, J., Qian, G., Chen, J., and Chen, S. (2012). FastUniq: A Fast De Novo Duplicates Removal Tool for Paired Short Reads. PLoS One 7, e52249.

Yang, Y., Torchinsky, M.B., Gobert, M., Xiong, H., Xu, M., Linehan, J.L., Alonzo, F., Ng, C., Chen, A., Lin, X., et al. (2014). Focused specificity of intestinal TH17 cells towards commensal bacterial antigens. Nature 510, 152–156.

Yin, Y., Wang, Y., Zhu, L., Liu, W., Liao, N., Jiang, M., Zhu, B., Yu, H.D., Xiang, C., and Wang, X. (2013). Comparative analysis of the distribution of segmented filamentous bacteria in humans, mice and chickens. ISME J. 7, 615–621.

Zimmermann, M., Zimmermann-Kogadeeva, M., Wegmann, R., and Goodman, A.L. (2019). Mapping human microbiome drug metabolism by gut bacteria and their genes. Nature.

