## Supplementary material for "Genome Sequence of Segmented Filamentous Bacteria Present in the Human Intestine"

.

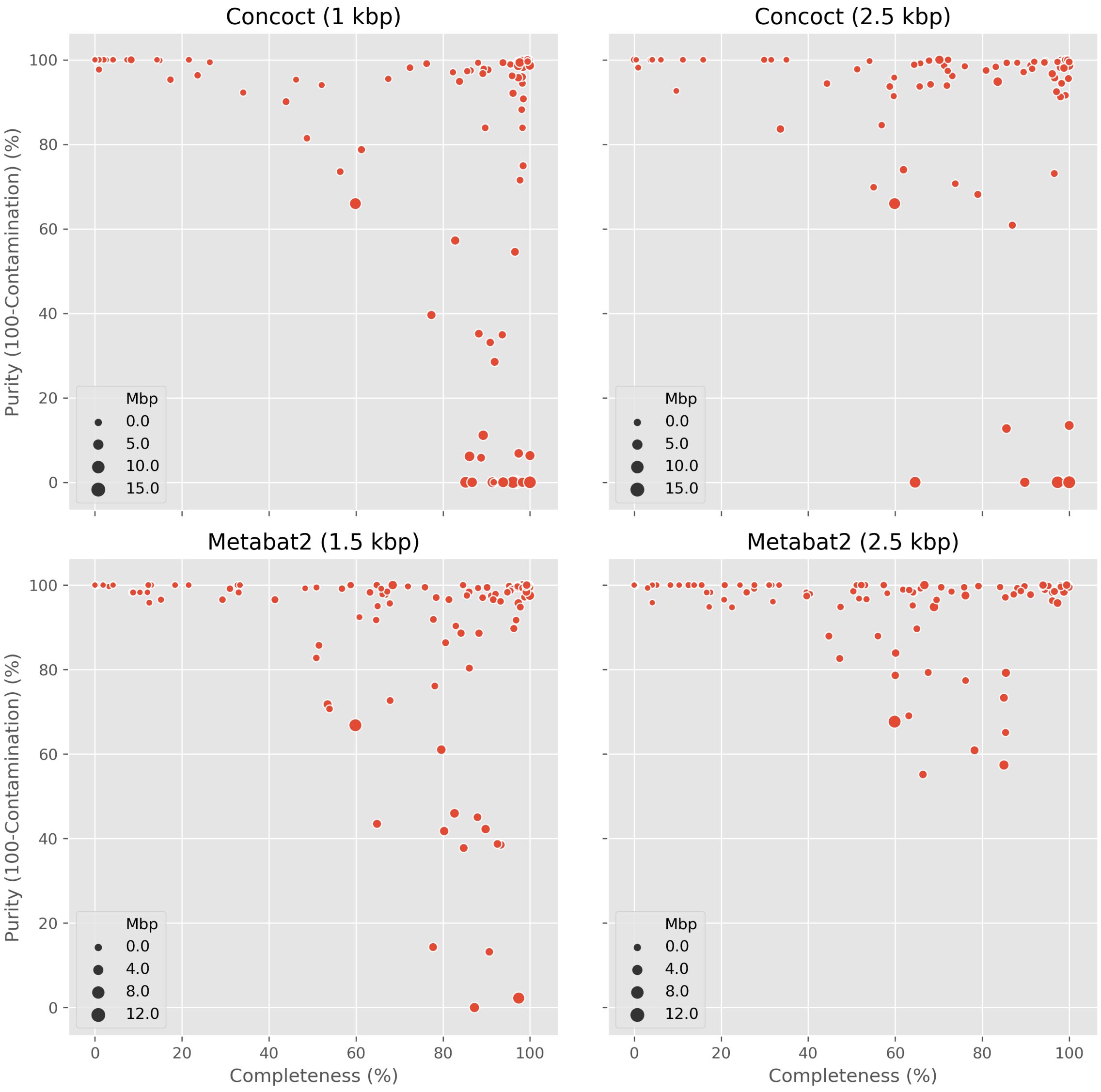


**Figure S1. Metagenomic bins obtained with CONCOCT and Metabat2 using different cutoffs on minimum contig length.** Each circle represents one bin, with its position along the x- and y-axis indicating estimated completeness and purity, respectively.

>sfb.merged_k141_89555_16S-rRNA
GATAAGAGTTTGATCCTGGCTCAGGACGAACGCTGGCGGCGTGCCTAACACATGCAAGTTGAACG
GAGTTATGTGGATTACTTTAGGGTAATTTATGTAATTTAGTGGCGAACGGGTGAGTAACACGTAG
ATAATCTGTCCTATATTGGGGGATAGGCCGATGAAAGTTGGATTAATACCGCATATAGCTAATAT
ATTGCATGATATATTAGTGAAAGGGAGAAATTTTGATATAGGAGGAGTCTGCGACACATTAGCTA
GTAGGTAAGGTAAAGGCTTACCTAGGCGACGATGTGTAGCTGGTCTGAGAGGATGGACGGCCACA
ATGGAACTGAGACACGGTCCATACTCCTACGGGAGGCAGCAGTGGGGAATATTGCACAATGGGGG
GAACCCTGATGCAGCAACGCCGCGTGAGTGAAGAAGGTTTTCGGATTGTAAAGCTCTGTTAGCAG
GGAAGAGGAAGGACGGTACCTGCAGAGGAAGCCACGGCTAACTACGTGCCAGCAGCCGCGGTAAT
ACGTAGGTGGCAAGCGTTGTTCGGAATAACTGGGCGTAAAGGATGCGTAGGCGGTTATATAAGTT
ATGTTGTTAAATATACAGGCTTAACCTGTAGAAAGCGGATAAAACTGTATGACTTGAGTGCAGGA
GAGGTAAGTGGAATTTCTAGTGTAGCGGTGAAATGCGTAGAGATTAGGAAGAACACCAGTGGCGA
AGGCGACTTACTGGACTGTAACTGACGCTGAGGCATGAGAGCATGGGGAGCAAACAGGATTAGAT
ACCCTGGTAGTCCATGCTGTAAACGATGGGTACTAGGTGTGGGTTGTGAATAGCAATCTGTGCCG
TCGCAAACGCAATAAGTACCCCGCCTGAGGAGTACGATCGCAAGATTAAAACTCAAAGGAATTGA
CGGGGACCCGCACAAGCAGCGGAGCATGTGGTTTAATTCGAAGCAACGCGAAGAACCTTACCTAG
ACTTGACATACCTTGAATTACCTTGTAATGAGGGAAGCTCGCAAGAGCAAGGATACAGGTGGTGC
ATGGTTGTCGTCAGCTCGTGTCGTGAGATGTTGGGTTAAGTCCCGCAACGAGCGCAACCCTTGTT
GTTAATTGCTAGCAGGTGAAGCTGAGCACTTTAGCGAGACAGCCTAGGTTAACTAGGAGGAAGGT
GGGGATGACGTCAAATCATCATGCCCCTTACGTCTAGGGCTACACACGTGCTACAATGGTGAGAA
CAGAGAGAAGCAAGCTAGTGATAGTGAGCAAACCTTATAAAACTCATCTCAGTTCGGATTGCAGG
CTGAAACTCGCCTGTATGAAGATGGAGTTGCTAGTAATCGCGAATCAGAATGTCGCGGTGAATAC
GTTCCCGGGTCTTGTACACACCGCCCGTCACACCATGAGAGTTGGCAACACCCGAAGCCTGTGAG
CTAACCGAAAGGAGGCAGCAGTCTAAGGTGGGGTTAATGATTGGGGTGAAGTCGTAACAAGGTAG
CCGTAGGAGAACCTGCGGCTGGATCACCTCCTTTC

Published SFB probes / PCR primers

| | | |
|---|---|---|
| SFBf1 | 5'-GGAGTCTGCGGCACATTAG-3' | Johnsson, 2013 |
| 779F | 5'-TGTGGGTTGTGAATAACAAT-3' | Urdaci, 2001 |
| 1008R | 5′-GCGAGCTTCCCTCATTACAAGG-3' | Snel, 1994 |
| 1008R | 5'-GCGGGCTTCCCTCATTACAAGG-3' | *Yin, 2013 |
| 1380R | 5'-GGTTAGCCCACAGGTTTCGG-3' | Urdaci, 2001 |
| 1380R | 5'-GGTTAGCCCACAGGCTTCGG-3' | *Yin, 2013 |
| SFBr1 | 5'-CACCTTAGACTGCTGCCTC-3' | Johnsson, 2013 |

*Sequences as given in Supplementary Table S2 in Yin et al.

**Figure S2. The 16S rRNA gene sequence of SFB-human-IMAG and published probes/primers used for the identification of SFB.** Nucleotides in red indicate mismatches relative to primers/probes. The Johnson and Yin references are given in the main article, Snel refers to Snel et al. System. Appl. Microbiol. 17, 172-179 (1994) and Urdaci to Urdaci et al. Res Microbiol 152: 67–73 (2001).

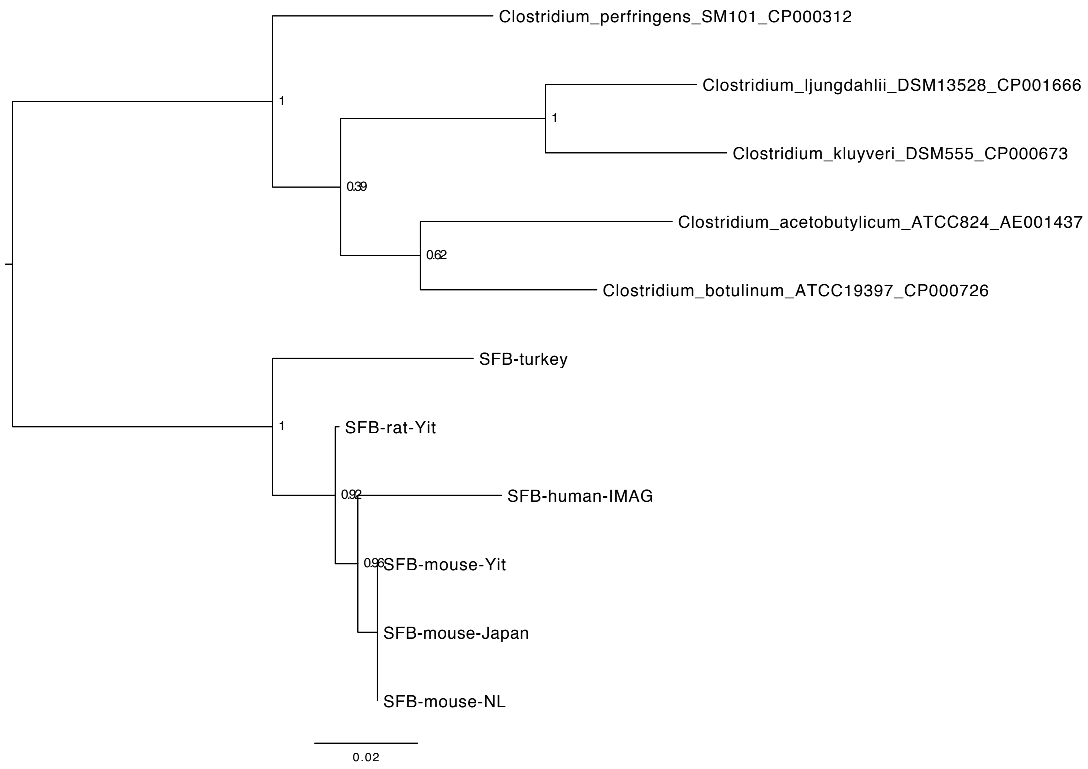


**Figure S3. Phylogenetic tree based on 16S rRNA gene sequences from sequenced SFB genomes.** The 16S rRNA gene sequences from five *Clostridium* genomes served as outgroup for rooting the tree.

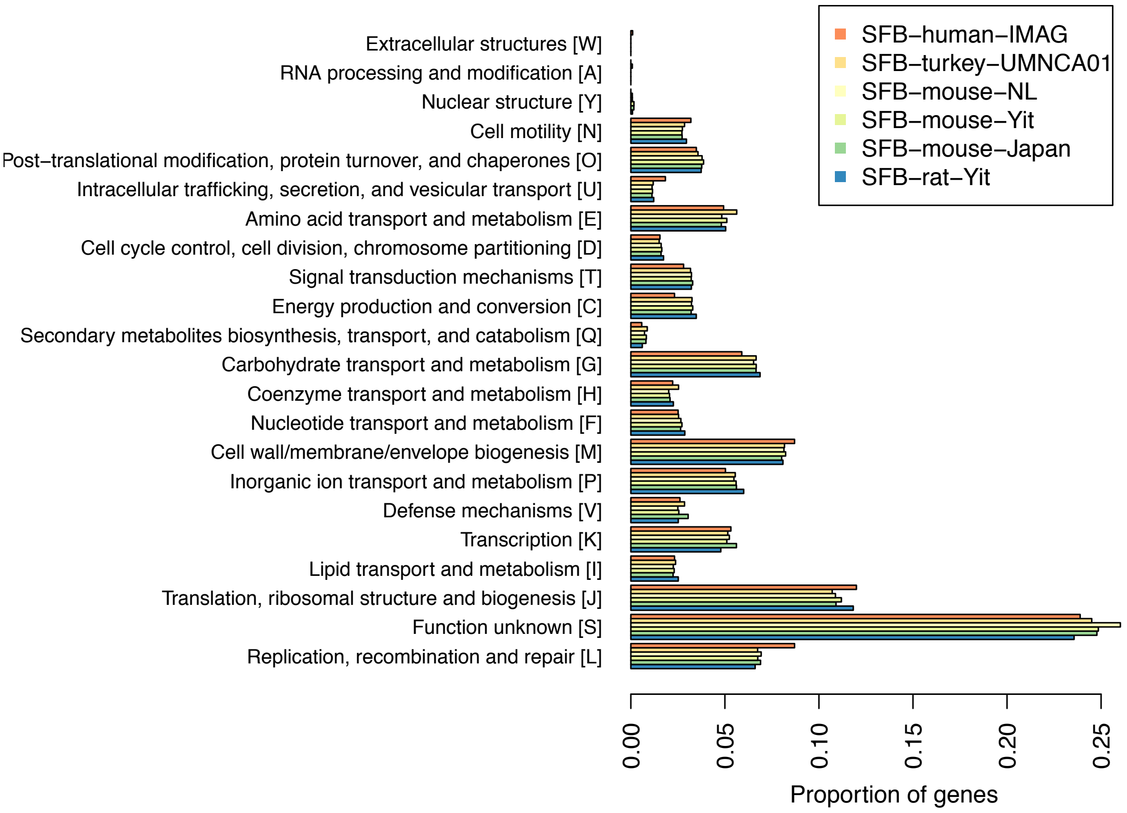


**Figure S4: Distribution of eggNOG functional categories in different SFB genomes.** The proportions were calculated based on genes having an eggnog annotation.

```
SFBM_0962           MKLTFNSNAMKTFMTYKKTISDHSKSIKNISTGKQINSAKDNPSRISKLANFEKEIRGYQ    60
00941               MKLNFNSNAVKTLRAYKKNILDHSKNINNISTGKKINSSKDNPNHISKLGNFEREIRGYQ    60
00379               MIINHNLNVMNAHRNMSNSVKSQGKSMEKLSSGLRIGRAGDDAAGLAISEKMRTQIRGLD    60
SFBM_0642           MIINHNINAMNAHRNMGSTTIAQGKAMEKLNSGLRINRAADDAAGFAISEKMRSQIRGLN    60
00424               MIINHNLNAMNAHRNMTGTTTAQGKAMEKLSSGLRINRAGDDAAGLAISEKMRSQIRGLD    60
SFBM_0583           MIINHNMNAMNAHRNMGSTTVSQGKSMEKLSSGLRINRAGDDAAGLAISEKMRSQIRGLN    60
SFBM_0584           MIINHNMNAMNAHRNMGSTTIAQGKAMEKLNSGLRINRAADDAAGLAISEKMRSQIRGLN    60
                    * :..* *.:::      .   :.* ::::.:* :*. : *:   ::   ::. :*** :

SFBM_0962           SSRRNIQDTVSMIQSADSVMGSLNDRVTRLKEITISLGNGSIQGDEKDIIQNEVNSILEG    120
00941               ASRRNIQDTVSMIQAADGVLDSVGNRITRIRELSVELGNASIGEDEKNILGNEINTLLEG    120
00379               QASRNAQDGIPMVQTAEGALTETHAITQRMRELAVQSANGTYTDEDREIINQEFTQLKKE    120
SFBM_0642           QASRNTQDGISMIQTAEGALSETQAIAQRMRELAVQSANGTYTDEDRTLIDQEFKKLKSE    120
00424               QASRNAQDGISMIQTAEGALSETQAIGQRMRELAVQSANGTYTDEDRKLIDQEFNQLKSE    120
SFBM_0583           QASRNAQDGISMIQTAEGALSETQAIGQRMRELAVQSANGTYTDEDRELINQEFNQLKSE    120
SFBM_0584           QASRNAQDGISMIQTAEGALSETQAIAQRMRELAVQSANGTYTDEDRTLIDQEFNQLKSE    120
                     : ** ** : *:*:*:..: .       *::*:::. .*.:   ::: :: :*.. : .

SFBM_0962           MSYDIEHFSFNEINILGNKDVTDNSNPKLI------------------------------    150
00941               IDYEIKNFSFNGINVLGHGETNDNSNPQLV------------------------------    150
00379               IDRIADDTDFGGVKVLSGSISSKNIQPGDVKRD----NGKCTMN----FYATGFKLTKQD    172
SFBM_0642           IDRISEDTEFNGSKLINGSISGKEIKTRDKDS--VII--NGDIKNPNSINEVMKEIDPNE    176
00424               IDRIATDTEFNGNKVINGNLSSKKIVEGT------VTKGTGSMNNVTEADLEFFEGDV--    172
SFBM_0583           IDRIANDTEFNGSKVLNGDKSGEKIEFDTGSITSTAVNGGTPLDA-ANAVNVFLETDLKN    179
SFBM_0584           IDRIATDTDFSGSKVLNGEKSGEKIEFESGTVASTG---TGTLNN-TDAAAVFLKTDLEN    176
                    :.    . .*.  :::.     .:

SFBM_0962           -----------------------SVLTT--------------------------------    155
00941               -----------------------EVLTT--------------------------------    155
00379               AKDLGEGNFTFKVSVSDDSKVTIDLIDKSYFNDYKEYIIDSYTFDGTDPSSTLGSRDMIK    232
SFBM_0642           LVRLGEGTFDLIISKDRNLDVTISIVDRSHFNGWKPYVIDSIIIDNPSVIEDVE------    230
00424               ANILGEGTFTLDIKIDAGGVININLLDKSRFNGNEEYVMDSITIDDTQIKADIGSANKQG    232
SFBM_0583           VNTLGEGTFHLKLTVDG-NEAKIELTDTSKFNDNKAYVLDSIVIKDGAVGNDEH------    232
SFBM_0584           VNTLGEGTFKLNLSLDG-DDVKLELIDESKFNDNKAYVLDSTVIKDGAADNDEH------    229
                                           .:

SFBM_0962           --------------------------------------------------GNSGEFSEIP    165
00941               --------------------------------------------------SKSGEVTEIP    165
00379               IKIDKAEINMGLT-------GLKPGGVA--YFDLVNASPPRGDGVILQVEANQNQRVGFS    283
SFBM_0642           ----LAGINLRFNNIHATSSYLPAG--KEIRISFFNDSEKKSNGVDFQVGSNKGQMVGLD    284
00424                AGFTIGGVDFVLDVE-----HLEKGKDIKLTITNTTEEKETGAGVELQIGANQNQMIGLS    287
SFBM_0583           -EITLGGADFKLAKLK---THIQDANVETVDLTLNNKVTEKTDGVELQVGANKDQMIGVS    288
SFBM_0584           -EITLGGAAFKLAQLK---DKIKDANVETVELTLNNVVKEKTNGVELQVGANKDQMISIS    285
                                                                  .:..:   .

SFBM_0962           SFNLSIENLGIDGMNV--KDDIGFNLEKIDNATSQIINARTKLGAISATFEDKIEQSESI    223
00941               LFNLSIESLGIKDFDV--NDDLSSILGKVDNAFSEVVSARSKLGAIGNTLEDKIKHSENL    223
00379               IENMSHRELGLGGINISTAENAQEAIARLDFSINAVSTQRADLGAIQNRLETIVTSTNNT    343
SFBM_0642           IENMRSRELGLGGSDLSTIENSKEAIERLDIAIFRVSEQRANLGSVQNRLEHTVSSIGNT    344
00424               MENMRSRELGLGGVNVATAESSQDAIGKLDEAISRISAQRADLGAAQNRLEHTIASTDNT    347
SFBM_0583           IGNMGSRELGLGGVNVSTAEDAKDAIGRLDEAVSRISAQRADLGAAQNRLEHTIASTDNT    348
SFBM_0584           IGNMRSRELGLGGVNVATTDDAKDAIGRLDEAVSRISAQRADLGASQNRLEHTIASTDNT    345
                      *:  ..**: . ::   :.    : ::* :   :   *:.**:    :*  :    .

SFBM_0962           EEVVTGAKSKIEDADVALEMLEFSRTLMLTEANIKNMSKTIYLPNDLIDVLGKLYK        279
00941               EEAIVGSKSKIEDSDIA---------------------------------------        240
00379               AENLQAAESRIRDVDMAKEMMNLTKLNILQQVS-----------------------        376
SFBM_0642           AENLQAAESRIRDVDMAKEMMNLTKLNILQQATQSMLAQANQSPQQVISILK----        396
00424               AENLQAAESRIRDVDMAKEMMNLTKLNVLQQASQSMLAQANQAPQQVLSILR----        399
SFBM_0583           AENLQAAESRIRDVDMAKEMMNLTKLNVLQQASQSMLAQANQAPQQVLSILR----        400
SFBM_0584           AENLQAAESRIRDVDMAKEMMNLTKLNVLQQASQAMLAQANQAPQQVLSLLK----        397
                     * : .::*:*.* *:*
```

**Figure S5. Clustal W alignment of the amino acid sequences of putative flagellins in SFB-Human-IMAG (00379, 00424, 00941) and SFM-mouse-Japan (SFBM_0583, SFBM_0584, SFBM_0642, SFBM_0962; Kuwahara et. al 2011)**. The yellow labeled amino acids depicts the most important positions for TLR5 signaling activity according to Song et. al 2017. Red color represents deviation from the consensus sequence.

```
00774              MNKKRIALALSAALGINTLMITVGNINGQVQIVHAVTDTRLND-SSIIDTLKVKTLKVQT     59
SFBNYU_003340      MNKKRVALALAAALGMNSLMVTVGAVGGQAFIAHAQDSKLSTDLLDKLKTLEIESLETSV     60
                   *****:****:****:*:**:*** :.**. *.**  ..  .*  . :.**::::*:...

00774              SMSSDGKDLYLSFPDRTDISNAKISSFNRQNFTERGNIDGNSGQVPATSSWITSGNYLSI     119
SFBNYU_003340      DMSTDGKAQVTFTKISPETISNVTGEFEYKPFTERSDLKETA--SKATGVWDKSTGKFNV     118
                   .**:***           :  .   ..*: : ****.::. .:    **. * .* . :.:

00774              SGFNSNAPGIYTTEVTITTNSGATETYKISLAKTPTDTLTAGKVT-VEPMSVTVEDLSLN     178
SFBNYU_003340      SGI--TAPGIYTGEVTIAYANKMTAKYTLSLVKKPSDSITFNTEVGVGT--VKLSNIEFD     174
                   **:  .****** ****:  .  * .*.:**.*.*:*::* .. . *    *.:.::.::

00774              GK-----ALDIGKKVFLLNSSDQEVSYGKVEDKAGTVKFNMSGVNM--KVGE----VYKI     227
SFBNYU_003340      NGSSKVTGFKPKKVSFSLNGGT-PVEVG--PDANGNYVLNLLDLGNKGNLKDTDKLTANF     231
                   .      .:.  *  * **..   *. *   *  *.  :*: .:.   :: :    . ::

00774              VYGVDKEDASALYKASTYLVMTRADNRGYDIVGATNVKDALSTAKAEIAKDFGVNTDSVA     287
SFBNYU_003340      SYGAN-----SEYTLSSDIVLVRKNQKITN-AVASNDKTKFPNIKQQIKTDFGLTTDPEI     285
                    **.:     : *. *: :*:.* :::  : . *:* *  : . * :* .***:.**

00774              IKETNTGYSVSVPKLNSSFDAKEETVLELPQTFGAASVLHGLPLTVTTVGNSLKVDFGDG     347
SFBNYU_003340      GT-----YVDNTHSYTISFGGK--ELLTYDQTVSTGDKLGGASLTLTTTGSQVKLSPESA     338
                    .     *  .. . . **..*   :*   **..:.. * *  **:**.*..:*:.  ..

00774              KI----TDTNNYVPYAQLSYGNNQS-----ETTLVGDYTATFNSGSTTLTTVSYAEFSNV     398
SFBNYU_003340      TLAKAAQLDDNYVPNVMLSYGINTAGNSTISAGLIGDYTLTIDKNTITCL-------DAM     391
                   .:       :**** . **** * :     .: *:**** *::..: *         . :

00774              TELKPSLTAVTNKA--------GTVSLGKKYDLKT---DDYTNSSANLDLSSFEHGCHEI     447
SFBNYU_003340      NGLTPSITTTNIANIDYSYVAFTGTGIGGNSELVTDMTKIDKNQSITLGIDKFEHGYNAI     451
                   . *.**:*:..              ..:* : :* *   .  .*.* .*.:..**** : *

00774              SIDFGVSVAADFATQVNDAAKNASSATASTITPNTEAKIKQSDDKSGDSVKTSILLAIDD     507
SFBNYU_003340      PVTFEVDAKNDKNTISA--------LTASMTGANNVIADINTASTSEEAVKASIIVSVDL     503
                    : * *..  *  *            ***    *.     :: ..* ::**:**::::*

00774              IDGVAVSSQFIQTDATKGKLLIKNGKSLFANGAPDSDIGS-KLSVSGAKS-VALSSSDTE     565
SFBNYU_003340      FDGVAVQSKFVKTSSSEGQVTIVGGSKLLNDLVGSLDYNEAVLTIDGVKATVNSTKSNDN     563
                   :*****.*:*::*.:::*:: * .*..*: : . . * ..  *::.*.*: *  :.*: :

00774              SLVYDVEFNNAVSDT--ISWELSGTSK-VKGET-AVQGKVAAANLSSVGIQN--APNTVT     619
SFBNYU_003340      DLIFDVQFNGAVPNKTDASWVLDIKGKKVEGKVDLTPGAVESVNLTSVQATNSGAENITD     623
                   .*::**:**.** :.   ** *. ..* *:*:.  . * * :.**:**   *  * * .

00774              KVKVNAQFTSDIPGLTAIGFGTQNTLEVDVNELTSTGAKSVRGSVSSGKYQGTYHAGVLV     679
SFBNYU_003340      QFKIDAKFNSTVPTGTVRVNDPVNGSDRVTFDQFKSGNNTIHNVVTSGKYQGTYSAGVIA     683
                   :.*::*:*.* :*  *.   .  *  :  . :  .:* ::::. *:******** ***:.

00774              DKVVGKITLEKPVTVTGNSVTLRFYTDFINEDDQNVNITKVNRSVIAYRRT----DGIGG     735
SFBNYU_003340      NYNPITVLLKA-DNPTTSSVRLTIDMDFFGNNDLIKNI---TGGIIYYREYKAGLGANDG     739
                   :    .: *:   . * .** * :  **:.::*   **   . .:* **.     .. .*

00774              WNITELSMEEAVNSITKTTTKTITGLERDTNYNFVVFMDVILTGGGFNGETHILQSNVVT     795
SFBNYU_003340      WEIAKGADISSSDIKEDPVTKVVTGLNAGKEYEFVVTYSYKEGN----SEIRAIYSNIAV     795
                   *:*:: :  .: :   . .**.:***: ..:*:***  .    .    .* : : **:..

00774              QRTNVNSSN-NNNNITGNGSNSTSGSSTGTTSIVVGNNNSTKTPTSLTLTLPGS-GY-ST     852
SFBNYU_003340      ATTKPGSSSSSGGTISGG---SSSSSSTGSTTVNVTSTNSTKGSTSISVTIPSGVSYTES     852
                     *: .**. ....*:*.   *:*.****:*:: * ..****  **:::*:*.. .* .:

00774              SYAPVVRGIKYKGTDGKVVTATSEQFANVKASYSSDGKSIILEGLVPDKDYTEISVDYTD     912
SFBNYU_003340      SANPTPVGFKYKDKNGKTVIETNDQYSNVKARFQN--GKVVLEGLVPGKEYTEISINYND     910
                   *  *.  *:***..:**.*  *.:*::**** :..   .::******.*:*****::*.*

00774              NNNKTRTLVIRDFKLTSVVDTLESYLRNVYTVVFDRPADEAGYNYHLNNLKSKKVSLRDF     972
SFBNYU_003340      KNNKTKTLVLKNVKIDSTS-EGQSYLANVYNVVFNRPADENGYMFHLGNLTSKKVSLRAF     969
                   :****:***:::.*: *.     :*** ***.***:***** ** :**.**.******* *

00774              LLNILNEKEFLGRYTTTEGKVEALYSAIVSRKSDEAGKAFWVNEYKTLLAVYGSEDNTLK     1032
SFBNYU_003340      LLNMINEKEFDKLYTTAENKVEALYNAIVNRDSDTQGKAFWVAEYKKLLSVYGSSSATLQ     1029
                   ***::*****   ***:*.******.***.*.**  ****** ***.**:****.. **:

00774              AIADRMVNENELKELANKLGVQW          1055
SFBNYU_003340      ALADRMVNENELKTLAQKLNFEW          1052
                   *:*********** **:**..:*
```

**Figure S6. Clustal W alignment of the amino acid sequences of gene 00774 from SFB-Human-IMAG and SFBNYU_003340.** Turquoise labelling indicates the predicted signal peptide. Yellow labelling indicates the $T_h17$ stimulating epitopes from SFBNYU_003340 as described earlier (Yang et al. 2017). Residues labelled in green depicts conserved positions in the C-terminally located motifs CR-1A, CR-2A, CR-2A and CR-2B earlier identified in many tentatively extracellular SFB proteins (Pamp et al. 2012). Histidine (H), colored in grey indicates deviation from the consensus motif described by Pamp et al.

| Parameter | Value |
|---|---|
| Completeness (%) | 85.58 |
| Contamination (%) | 0 |
| Genome size (bp) | 1314549 |
| Number of contigs | 153 |
| N50 contig length (bp) | 11763 |
| Mean contig length (bp) | 8591 |
| Longest contig (bp) | 55026 |
| GC (%) | 26.98 |
| Number of protein-coding genes | 1276 |
| Number of 16S rRNA genes | 1 |
| Number of 28S rRNA genes | 1 |
| Number of tRNA genes | 28 |

**Table S1. Summary statistics for SFB-human-IMAG.**

| COG_ID | COG_category | SFB-human-IMAG_gene(s) | turkey-UMNCA01 | human-IMAG |
|---|---|---|---|---|
| COG0208 | F | sfb.merged_00330 | 0 | 1 |
| COG1760 | E | sfb.merged_00449<br>sfb.merged_00452 | 2 | 2 |
| COG1937 | S | sfb.merged_00457 | 0 | 1 |
| COG0607 | P | sfb.merged_00458<br>sfb.merged_00459 | 0 | 2 |
| COG1424 | H | sfb.merged_00527 | 1 | 1 |
| COG0132 | H | sfb.merged_00530 | 1 | 1 |
| COG0161 | E | sfb.merged_00531 | 1 | 1 |
| COG0156 | H | sfb.merged_00528<br>sfb.merged_00756<br>sfb.merged_00757 | 2 | 3 |
| COG2188 | K | sfb.merged_00523<br>sfb.merged_00973 | 1 | 2 |
| COG1887 | M | sfb.merged_00005<br>sfb.merged_00006 | 0 | 2 |
| COG1514 | J | sfb.merged_00007 | 0 | 1 |
| COG4912 | L | sfb.merged_00584 | 1 | 1 |
| COG1715 | V | sfb.merged_00719 | 0 | 1 |
| COG0451 | M | sfb.merged_00758 | 1 | 1 |
| COG2189 | L | sfb.merged_00768<br>sfb.merged_00769<br>sfb.merged_00770 | 0 | 3 |
| COG0300 | S | sfb.merged_00793 | 0 | 1 |
| COG3505 | U | sfb.merged_00808 | 0 | 1 |
| COG3451 | U | sfb.merged_00813<br>sfb.merged_00814 | 0 | 2 |
| COG3942 | S | sfb.merged_00816 | 0 | 1 |
| COG0827 | L | sfb.merged_00819<br>sfb.merged_00820<br>sfb.merged_00821<br>sfb.merged_00822 | 0 | 4 |
| COG1670 | J | sfb.merged_00949 | 2 | 1 |
| COG1321 | K | sfb.merged_00961 | 1 | 1 |
| COG1680 | V | sfb.merged_01000 | 0 | 1 |
| COG1737 | K | sfb.merged_01007 | 2 | 1 |
| COG3862 | S | sfb.merged_01065 | 1 | 1 |
| COG1313 | S | sfb.merged_01287 | 0 | 1 |
| COG0517 | S | sfb.merged_01293 | 0 | 1 |
| COG1278 | K | sfb.merged_01295 | 0 | 1 |
| COG3843 | U | sfb.merged_00120 | 0 | 1 |

**Table S2. COGs found in SFB-human-IMAG that are not found in any of the complete SFB genomes from rodents.** The integers indicate the number of times the COGs occur in the SFB genomes from turkey and human.

| CAZyme | SFB-human-IMAG | Secretion signal |
|---|---|---|
| CE4 | sfb.merged_00222 | SP |
| CE4 | sfb.merged_00371 | SP |
| CE4 | sfb.merged_00421 | SP |
| CE4 | sfb.merged_00862 | - |
| CE4 | sfb.merged_01243 | SP |
| GH1 | sfb.merged_00525 | - |
| GH144 | sfb.merged_00150 | - |
| GH23 | sfb.merged_00563 | SP |
| GH23 | sfb.merged_01271 | - |
| GH3 | sfb.merged_00630 | |
| GH89 | sfb.merged_00532 | SP |
| GH94 | sfb.merged_00096 | - |
| GH94 | sfb.merged_00149 | - |
| GH94 | sfb.merged_01168 | - |
| GT2 | sfb.merged_00325 | - |
| GT2 | sfb.merged_01223 | - |
| GT2 | sfb.merged_01225 | - |
| GT26 | sfb.merged_00218 | - |
| GT28 | sfb.merged_00181 | - |
| GT28 | sfb.merged_01254 | - |
| GT4 | sfb.merged_00220 | - |
| GT4 | sfb.merged_00863 | - |
| GT4 | sfb.merged_01184 | - |
| GT4 | sfb.merged_01185 | - |
| GT4 | sfb.merged_01277 | - |
| GT51 | sfb.merged_00191 | - |
| GT51 | sfb.merged_00331 | - |
| GT84 | sfb.merged_00096 | - |

**Table S3. Carbohydrate-active enzymes (CAZymes) in SFB-human-IMAG.** CAZyme anotations were inferred using hmmscan against the dbCAN (http://bcb.unl.edu/dbCAN/) database. Reported hits all have e-value < 1e-18 and coverage > 0.35. Secretion signal was predicted with SignalP.
